# NRF2 activation in the heart induces glucose metabolic reprogramming and mediates cardioprotection via upregulation of the pentose phosphate pathway

**DOI:** 10.1101/2023.05.12.540342

**Authors:** Anna Zoccarato, Ioannis Smyrnias, Christina M Reumiller, Anne D Hafstad, Mei Chong, Daniel A Richards, Celio XC Santos, Asjad Visnagri, Daniel I Bromage, Xiaohong Zhang, Greta Sawyer, Richard Thompson, Ajay M Shah

## Abstract

**Rationale:** The transcription factor NRF2 is well recognized as a master regulator of antioxidant responses and cytoprotective genes. Previous studies showed that NRF2 protects mouse hearts during chronic hemodynamic overload at least in part by reducing oxidative stress. Evidence from other tissues suggests that NRF2 may modulate glucose intermediary metabolism but whether NRF2 has such effects in the heart is unclear.

**Objective:** To investigate the role of NRF2 in regulating glucose intermediary metabolism and cardiac function during disease stress.

**Methods and Results:** Cardiomyocyte-specific Keap1 knockout (csKeap1KO) mice, deficient in the endogenous inhibitor of NRF2, were used as a novel model of constitutively active NRF2 signaling. Targeted metabolomics and isotopomer analysis were employed in studies with ^13^C_6-_glucose in csKeap1KO and wild-type (WT) mice. Pharmacological and genetic approaches were utilized in neonatal rat ventricular cardiomyocytes (NRVM) to explore molecular mechanisms. We found that cardiac-specific activation of NRF2 upregulated key enzymes of the Pentose Phosphate Pathway (PPP), redirected glucose metabolism towards the PPP and protected the heart against pressure overload-induced cardiac dysfunction. *In vitro*, knockdown of Keap1 upregulated PPP enzymes and reduced cell death in NRVM subjected to chronic neurohumoral stimulation. These pro-survival effects were abolished by pharmacological inhibition of the PPP or silencing of the PPP rate-limiting enzyme glucose-6-phosphate dehydrogenase (G6PD). Knockdown of NRF2 in NRVM increased stress-induced DNA damage which was rescued by supplementing the cells with either NADPH or nucleosides, the two main products of the PPP. Activation of NRF2 also protected the heart against myocardial infarction-induced DNA damage, dysfunction, and adverse remodeling.

**Conclusions:** These results indicate that NRF2 regulates cardiac metabolic reprogramming by stimulating the diversion of glucose into the PPP, thereby providing cardiac protection during stress by generating NADPH and providing nucleotides to prevent stress-induced DNA damage.

## INTRODUCTION

Heart failure is a leading cause of death worldwide and represents a substantial socioeconomic burden^1^. In response to chronic pathological stresses such as hemodynamic overload or myocardial infarction, the heart undergoes complex remodeling which in the long term may lead to heart failure. Pathological cardiac remodeling is characterized by the activation of multiple pathways that collectively drive structural and functional changes including contractile dysfunction, metabolic alterations, cardiomyocyte death, fibrosis, and ventricular dilatation^2, 3^. An increase in oxidative stress, resulting from an increase in reactive oxygen species production (ROS) and a decrease in cellular antioxidant capacity, is a universal feature of heart failure that impacts on diverse aspects of pathological cardiac remodeling^4, 5^.

Previous studies showed that the transcription factor nuclear factor erythroid–derived 2-like 2 (NRF2) is involved in the response to pathological cardiac stress. NRF2 functions as a key controller of the cellular redox status. Under physiological conditions, NRF2 levels are kept low by binding to the inhibitor protein, Kelch-like ECH–associated protein 1 (Keap1), which promotes NRF2 ubiquitination and proteasomal degradation. During cell stress, key functional Keap1 cysteines become oxidized thus weakening its binding to NRF2 and promoting NRF2 nuclear translocation and transcriptional activation of its target genes^6^. These include numerous antioxidant and detoxification enzymes, such as glutathione S-transferases (GSTs), NAD(P)H:quinone oxidoreductase-1 (NQO1), the glutamate–cysteine ligase catalytic (GCLC) and modifier (GCLM) subunits which catalyse the rate-limiting step in glutathione (GSH) biosynthesis, glutathione reductase (GSR) which reduces oxidized glutathione, and thioredoxin reductase-1 (TRXR1) which reduces oxidized protein thiols^7^. Our group and others have reported that NRF2 is essential in modulating adaptation and protection against cardiac injury and contractile dysfunction in response to pathological stresses, at least in part via activation of antioxidant pathways^8–12^. Aside from its role as a redox homeostasis regulator, NRF2 has recently been described to induce cellular metabolic reprogramming in stress conditions^13^. Evidence from the cancer field describes that NRF2, which in certain cancer types is associated with a malignant phenotype and cancer growth, redirects glucose and glutamine into anabolic pathways such as the pentose phosphate pathway (PPP) and purine biosynthesis to sustain cancer cell proliferation and survival^14^. Regulation of PPP enzymes including the NADPH-generating Glucose-6-phosphate dehydrogenase (G6PD) and 6-phosphogluconate dehydrogenase (PGD), and enzymes of the nonoxidative arm of PPP, transaldolase (TALDO1) and transketolase (TKT), by NRF2 has been reported in cancer cells^13, 14^ and other tissues including inflammatory cells^15^, liver^16^, primary cortical astrocytes^17^, and stem cells^18^.

In the heart, changes in energy substrate utilization and metabolism are key features of stress-induced cardiac hypertrophic remodelling^19–21^. Interesting parallels have recently been drawn between cancer cells and failing hearts in terms of metabolic reprogramming, which may allow for increased fluxes into intermediary metabolic pathways that help cells cope with the augmented energy and biosynthetic requirements^22^. In fact, chronic haemodynamic stress in the heart is associated with decreased fatty acid oxidation and increased glucose utilisation^19, 20^. Some recent studies have explored the role of glycolytic branch pathways in settings of pathological remodeling although with contrasting results^21, 23^. However, whether NRF2 activation regulates glucose metabolic reprogramming in the heart under pathological stress conditions has not yet been explored. In this study, we investigate the role of NRF2 in regulating cardiac metabolism and describe the effect of constitutive activation of NRF2 on cardiac glucose metabolism using state-of-the-art metabolomics. We show that activation of NRF2 increase glycolysis and glucose flux into the PPP, as well as the expression levels of key PPP regulating enzymes. Activation of NRF2 protects the heart from adverse remodeling and cardiac dysfunction upon chronic pressure-overload or myocardial infarction and reduces cardiomyocyte death, at least in part via PPP-mediated reduction of cardiomyocyte DNA damage.

## MATERIALS AND METHODS

### Mouse models

All animal procedures in this study were carried out according to the Guidance on the Operation of the Animals (Scientific Procedures) Act, 1986 (United Kingdom). Cardiomyocyte-targeted knockout of Keap1 was achieved by crossing Keap1 F/F mice^24^ with cardiomyocyte specific Cre (αMHC-Cre) mice, all on a C57BL/6 background. Aortic constriction was performed on male csKeap1KO mice and wild-type littermates (16-18g) by a single surgeon, using suprarenal banding with a 27-gauge constriction under 2% isoflurane/98% oxygen anaesthesia^25^. Sham constriction involved identical surgery apart from band placement. Animals were studied up to 6 weeks post-surgery. Female csKeap1KO mice and WT littermates approximately 12 weeks of age underwent left anterior descending coronary artery (LAD) ligation under 2% isoflurane/98% oxygen anaesthesia^26^. Sham groups underwent identical surgery except for ligation. Experimental analyses were done at 4 weeks after surgery.

### Echocardiography

Animal were imaged^27^ under 1.5% isoflurane using a Visualsonics Vevo 2100 ultrasound system with a 40-MHz transducer. Analysis of data was performed using the VevoStrain software package (Visualsonics).

### Langendorff Heart Perfusion

The preparation and perfusion of the hearts was performed in the Langendorff mode. In brief, mice were anaesthetized with an i.p. injection of pentobarbital sodium (90 mg/ kg) containing heparin (5000 U/ kg). The hearts were excised and cannulated and subsequently perfused with Krebs-Henseleit (KH) bicarbonate buffer containing 5 mmol/L glucose, 0.2 mmol/L pyruvate, 0.5 mmol/L glutamine, 1.5 mmol/L lactate, 0.4 mmol/L Na-octanoate, 50 mU/L insulin, 118.5 mmol/L NaCl; 5.9 mmol/L KCl; 0.48 mmol/L EDTA; 1.16 mmol/L MgSO_4_; 2.2 mmol/L CaCl_2_; 25 mmol/L NaHCO_3_ (37°C, pH 7.2). After 20 min equilibration, hearts were perfused for 20 minutes with a modified KH-buffer containing uniformly labelled glucose (5 mmol/L [U-^13^C]glucose, Cambridge Isotope Laboratories, Inc.) instead of standard glucose, with all other components unchanged. The perfusion pressure was set to 60 mmHg and monitored using an intraventricular balloon. Immediately after perfusion, hearts were freeze-clamped for metabolic quenching and stored at −80 °C.

### Metabolite extraction from tissue samples for LC/MS

Frozen heart tissue (∼40-50 mg) was lyophilized overnight (−55 °C) and subsequently ground in liquid nitrogen with a mortar and pestle. Pulverized tissue was then resuspended in a 4 °C cold chloroform: methanol solution (2:1, v/v) and vortexed. Resuspended tissues were incubated for 1 hour at 4 °C, including 3 pulses of sonication (8 min each). Subsequently, samples were centrifuged (0°C, 10 min, 16,000g), and the resulting supernatant was transferred to a new tube and dried at room temperature using a SpeedVac concentrator (ThermoFisher Scientific). The pellet was re-extracted by repeating this procedure with methanol: water solution (2:1, v/v). The dried metabolite extracts were reconstituted in chloroform: methanol: water solution (1:3:3, v/v/v) and incubated for 10 min before a final centrifugation step (0 °C, 5 min, 16,000g). After spinning, the upper polar phase was isolated and dried in a SpeedVac concentrator. Dried polar fractions were stored at −80 °C until MS analysis.

### LC-MS analysis

The LC-MS method has been previously described^28^. Briefly, the dried polar metabolite fractions were reconstituted in acetonitrile/water (v/v 3/2), vortexed and centrifuged at 16,000 g for 3 min before analysis on a 1290 Infinity II ultrahigh performance liquid chromatography (UHPLC) system coupled to a 6546-quadrupole time-of-flight (Q-TOF) mass spectrometer (Agilent Technologies). Samples were separated on a Poroshell 120 HILIC-Z column (100 × 2.1 mm, 2.7 μm, Agilent) attached to a guard column (5 × 2.1 mm, 2.7 µm) and analyzed in negative ionization mode using water with 10mmol/L ammonium acetate (solvent A) and acetonitrile with 10mmol/L ammonium acetate (solvent B), both solvents containing 5 µmol/L Infinity Lab deactivator additive (Agilent Technologies). The elution gradient used was as follows: isocratic step at 95% B for 2 min, 95 to 65% B in 12 min, maintained at 65% B for 3 min, then returned to initial conditions over 1 min, and then the column was equilibrated at initial conditions for 8 min. The flow rate was 0.25 mL/min; the injection volume was 1μL, and the column oven was maintained at 30 °C. Feature annotation and metabolite identification were based on accurate mass and standard retention times with a tolerance of ±5 ppm and ±0.5 min, respectively, and performed with MassHunter Profinder (version 10.0.2, Agilent Technologies) using our in-house curated metabolite library based on metabolite standards (Sigma-Aldrich). C^13^ label incorporation levels were normalized to the natural occurrence of C^13^ isotopes and are represented as corrected abundance percentages. Samples were run in one batch and injected into technical duplicates. Downstream pathway enrichment analysis was performed using Metaboanalyst 5.0 Software^29^.

### In vivo infusion of [U-^13^C]glucose

In vivo infusion of [U-^13^C]glucose was performed as previously described^30^. Briefly, mice were fasted for 6 h before glucose infusion. Animals under 1.5% isoflurane were given an intraperitoneal bolus of 0.4 mg/g (100 µL) [U-^13^C]glucose followed by a continuous tail vein infusion of 0.012 mg·g^−1^·min^−1^ at 150 µL/h for 30 min. Body temperature was maintained at 37°C using a thermostatic blanket, and respiratory rate was monitored using a MouseMonitor platform (Indus Instruments). At the end of the infusion, the heart was rapidly flushed with saline before being freeze-clamped in situ and then excised for storage in liquid N_2_.

### Metabolite extraction from heart samples and 1D NMR ^1^H-^13^C Heteronuclear Spin Echo

Metabolite extraction from heart samples was performed as previously described^30^. Briefly, heart tissue was minced before homogenization in an extraction buffer containing 1:1:1 methanol-chloroform-double-distilled H_2_O at a ratio of 1.7 mL extraction buffer per 0.2 g of sample. The tissue was homogenized in a Precellys homogenizer (Precellys) at 5,000 revolutions/min (rpm) for 2×20 s. The extract was vortexed for 15 min before centrifugation at 1,500 rpm at 4°C for 30 min. The supernatant was collected and dried under an Eppendorf dryer at 30°C. The final dried extracts were reconstituted in 170 µL of 100% deuterium oxide containing 0.5 mmol/L DSS, 100 mmol/L sodium phosphate (pH 7.0), and 6 mmol/L imidazole. NMR spectra were acquired at 25°C on a Bruker Avance 700-MHz spectrometer equipped with 5-mm triple-resonance z-axis gradient cryogenic probes. The ^13^C labeled metabolites were calculated using heteronuclear single quantum coherence (HSQC) as described previously^30^. Spectra were processed using the MetaboLab software package^31^.

### Histology

Hearts were arrested in diastole with 5% KCl and fixed with 2% paraformaldehyde either for 6 hrs at room temperature or overnight at 4°C. Subsequently, 6 µm paraffin-embedded transverse cross-sections were stained with FITC-conjugated wheat germ agglutinin (FITC-WGA, Vector RL-1022) to outline cardiomyocytes. Interstitial fibrosis was assessed by blinded quantitative image analysis (Volocity, Perkin Elmer) of Picrosirius red-stained sections ^10^. Apoptosis was assessed by terminal deoxynucleotidyl transferase dUTP nick-end labeling (TUNEL) staining following manufacturer’s instructions (Millipore S7110 kit). Infarct size after LAD ligation was measured by Evans blue dye (1%) staining as previously described^32^. To quantify DNA damage, sections were stained with an anti– γH2A.X antibody (Phospho-Histone H2A.X (Ser139) (20E3), Cell Signalling #9718). Imaging was done on a Nikon spinning disk confocal microscope.

### Western blotting

Heart tissue samples or pelleted cardiomyocytes were homogenized and lysed in hypotonic lysis buffer [50 mmol/L Hepes, pH 7.4, 10 mmol/L KCl, 5 mmol/L EDTA, 5 mmol/L EGTA, 2 mmol/L MgCl_2_, 2 mmol/L DTT, 0.1% NP-40, protease inhibitor cocktails and Ser/Thr and Tyr phosphatase inhibitor cocktails (Sigma, UK)]. Protein concentration for each sample was measured using BCA assay (Pierce, UK). Heart tissue homogenates or cell lysates were separated by SDS/PAGE and transferred onto nitrocellulose membranes. Antibodies used were the following: glucose 6-phosphate dehydrogenase (Abcam, ab210702), 6-phosphogluconate dehydrogenase (Abcam, ab129199), transaldolase 1 (ThermoFisher Scientific, A304-326A-T), transketolase (Cell Signalling, #8616), phospho-histone H2A.X (Ser139) Cell Signalling, #9718). Protein band quantification was undertaken using an Odyssey Li-Cor imaging system (Li-Cor Biosciences, UK)

### cDNA synthesis and real time qPCR

Total RNA was isolated from heart tissues or cardiomyocyte samples according to the manufacturer’s protocol (Qiagen). cDNA was synthesized using Oligo-dTs and M-MLV RT (Promega). qRT-PCR was performed with the StepOnePlus System (Applied Biosystems) using SYBR Green and the comparative Ct method was used, with cytoskeletal β-actin levels used for normalization. Primer sequences are listed in Table 1:

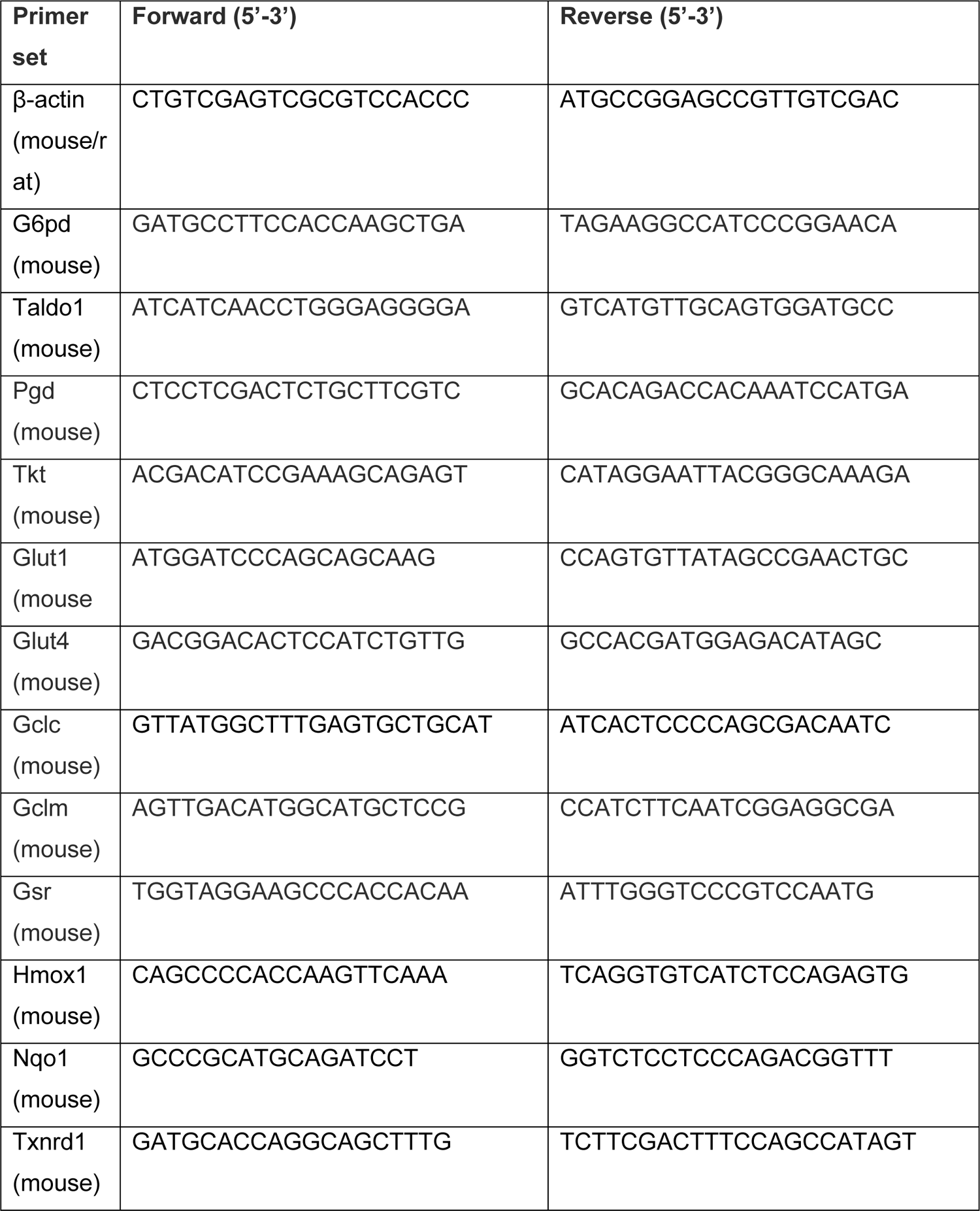
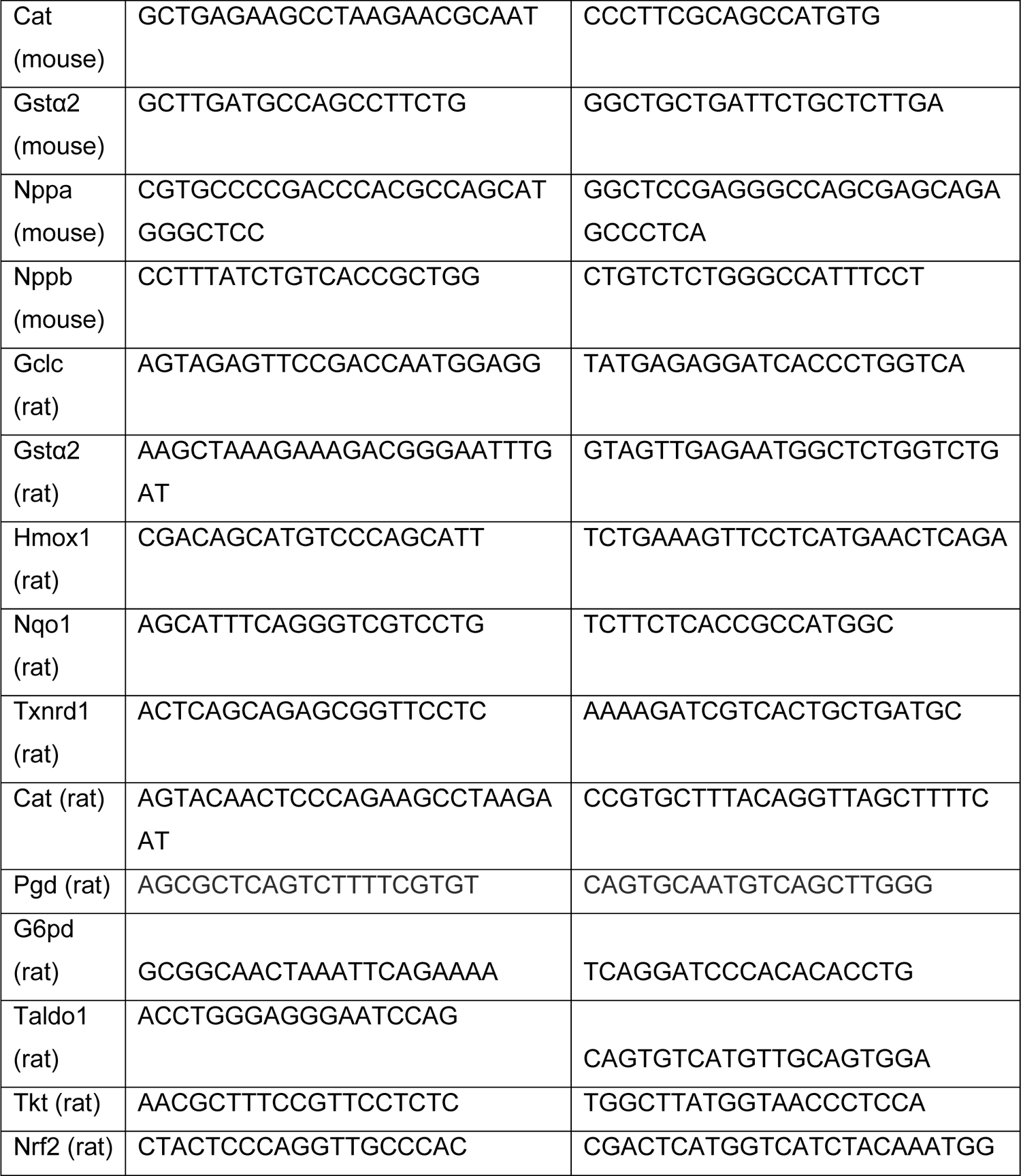

### Neonatal rat ventricular myocytes (NRVM)

Primary neonatal rat ventricular myocytes were isolated from 1 to 2-days old Sprague-Dawley rats (Charles River Laboratories) as previously described^33^. Cells were cultured in plating medium (DMEM High Glucose, MEM199, Horse serum, non-heat inactivated Fetal Bovine Serum, non-essential amino acids, Glutamine, Pen/Strep) for 24 hours before switching to a serum-free medium containing DMEM High Glucose, MEM199, Glutamine, Pen/Strep. Cells were transduced with adenoviral vectors (Welgen Inc.) expressing a short hairpin sequence targeted against NRF2 (shNRF2) or a non-targeting sequence (shCtr) at MOI 500. For the knockdown of Keap1 or G6pd, short interfering RNAs (siRNAs) were purchased from Thermo Fisher Scientific (Silencer® Select: s138759; Silencer® Select: s127750) and control siRNA was obtained from Dharmacon (ON-TARGETplus Non-targeting Pool). NRVM were transfected using TurboFect Transfection Reagent – (Thermo Fisher Scientific) following manufacturer’s instructions. NRVM were treated for 24 h or 8 h with isoproterenol 100µmol/L (Isoprenaline hydrochloride, I5627, Sigma-Aldrich), 6-aminonicotinamide 2.5mmol/L (A68203, Sigma-Aldrich), nucleosides 0.3 mmol/L (EmbryoMax® Nucleosides (100X), Merck) or NADPH 0.3 mmol/L (NADPH, Tetrasodium Salt, 481973, Merck).

### Immunofluorescence and confocal imaging

NRVM were washed with PBS and fixed in 4% PFA for 20 min at room temperature and permeabilized in 0.1% v/v Triton™ X-100 in PBS. Blocking and antibody incubations were performed in 2% w/v BSA. For cell viability assays 7-AAD (7-Aminoactinomycin D, Invitrogen, A1310) and Hoechst 33342 (Invitrogen, H3570) were used following the manufacturer’s instructions. Images were captured using a Nikon Spinning disk confocal and analysis was performed using ImageJ.

### Other assays

NADPH/NADP^+^ ratio was measured in NRVM using NADP/NADPH-Glo™ Assay (Promega, G9081) following manufacturer’s instructions.

### Statistical analysis

All data are represented as mean±SEM. Unpaired, 2-tailed Student’s t-test was used to compare differences between 2 groups. For multiple group comparisons with one variable, one-way ANOVA was performed, followed by Tukey multiple comparisons test. For multiple group comparisons with two or more variables, 2-way ANOVA was conducted, followed by Tukey multiple comparisons test. Number of replicates is indicated in the figure legends. Statistical analyses were performed with GraphPad Prism 9.5.1.

## RESULTS

### Cardiac-specific constitutive activation of NRF2 induces antioxidant responses and metabolic reprogramming in the heart

To study if NRF2 mediates metabolic reprogramming in the heart, we generated a mouse model of cardiac-specific constitutive activation of NRF2 by knocking-out Keap1 in cardiomyocytes. Conditional Keap1 floxed mice^24^ were crossed with cardiomyocyte specific Cre transgenic mice (αMHC [α-myosin heavy chain]-Cre) to generate cardiomyocyte-specific Keap1 knockout (csKeap1KO) mice (Supplementary Figure 1A). Because the Keap1 floxed model exhibits some hypomorphism^34^, we compared csKeap1KO with wild-type littermates (WT). csKeap1KO mice and WT littermates manifested no obvious differences at birth, had similar patterns of growth (Supplementary Figure 1B) and showed comparable baseline contractile function and cardiac dimensions by echocardiography (Supplementary Figure 1C-H). csKeap1KO hearts had significantly lower levels of Keap1 transcript and increased mRNA expression levels of NRF2 targets such as *Gsta2*, *Nqo1* and *Txnrd1*, confirming an activation of NRF2 (Figure 1A). To assess whether NRF2 activation in the heart mediates metabolic reprogramming, we performed targeted metabolomics encompassing glycolytic, branch pathway and TCA cycle metabolites, using liquid chromatography mass spectrometry (LC-MS) of WT and csKeap1KO hearts. Principal component analysis, performed using MetaboAnalyst 5.0, identified a clear separation and clustering between WT and csKeap1KO heart metabolite profiles (Figure 1B). The majority of the top 60 metabolites that differentially clustered between the two genotypes comprised metabolites related to the PPP, nucleotides and glutathione metabolism (Figure 1C). Indeed, csKeap1KO hearts had higher levels of glutathione (GSH and GSSG) and an increased GSH/GSSG ratio compared to WT littermates (Figure 1D). They also showed significantly higher mRNA expression levels of NRF2-regulated enzymes involved in glutathione synthesis and reduction such as *Gclc*, *Gclm* and *GSR* compared to WT littermates (Figure 1E). Overall, these data indicate that activation of NRF2 in the heart alters not only antioxidant mechanisms but also its metabolite profile.

**Fig. 1:**
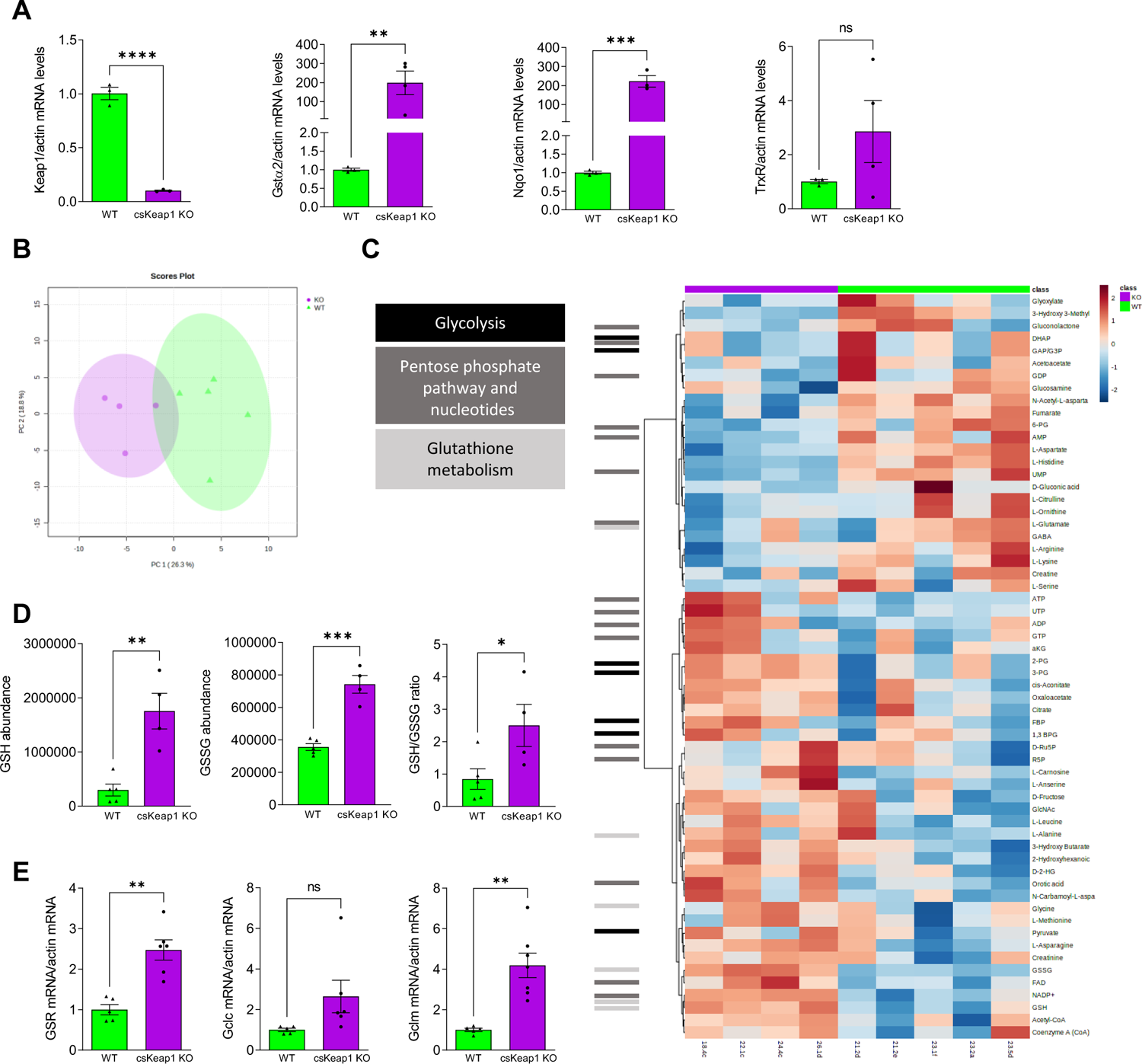
Cardiac-specific activation of NRF2 induces metabolic reprogramming in the heart. **A)** mRNA levels of the NRF2 targets glutathione-S-transferase α2 (*Gsta2*), nicotinamide adenine dinucleotide phosphate-quinone-oxidoreductase 1 (*Nqo1*) and thioredoxin reductase 1 (*Txnrd1*) in hearts of cardiomyocyte-specific Keap1 knockout (csKeap1KO) and wild-type (WT) control mice. n≥3/group. **B)** Principal component analysis of targeted metabolomic profile of csKeap1KO vs WT hearts, n≥4/group. **C)** Heatmap showing hierarchical clustering of metabolite abundance for the top 60 features in WT vs csKeap1KO hearts. **D)** Levels of reduced (GSH), oxidized (GSSG), and GSH/GSSG ratio in WT and csKeap1KO hearts measured by LC-MS, n≥4/group. **E)** mRNA expression levels of glutathione-disulfide reductase (*Gsr*), glutamate-cysteine ligase catalytic subunit (*Gclc*) and glutamate-cysteine ligase modifier subunit (*Gclm*) in WT vs csKeap1KO hearts, n≥5/group. Data are presented as mean ± SEM. **P < 0.01 and ***P < 0.001, ns, not significant by unpaired Student’s t test.

### NRF2 activation in the heart increases glucose utilization and flux into the PPP

Given the differences observed in abundance of metabolites related to glycolysis and the PPP, we assessed whether glucose utilization and flux were altered in the csKeap1 KO hearts. We undertook stable isotope-resolved metabolomics of *ex-vivo* csKeap1KO and WT Langendorff-perfused hearts using uniformly labelled glucose ([U-^13^C]glucose) followed by targeted LC-MS metabolomics and ^13^C-isotopomer analysis. A principal component analysis showed that csKeap1KO hearts presented a distinct ^13^C metabolite enrichment as compared to WT littermates (Figure 2A), with a large number of glycolytic metabolites showing differences in ^13^C enrichment between the two genotypes (Figure 2B). There was a significant increase in csKeap1KO hearts compared to WT in the % ^13^C label enrichment of glucose, glucose-6-phosphate (G6P), fructose-6 phosphate (F6P), phosphoenolpyruvate (PEP) and lactate (Figure 2C), and a trend towards an increase in ^13^C enrichment of PPP metabolites such as ribose-5 phosphate (R5P) and sedoheptulose-7 phosphate (S7P) (Figure 2E). No differences in ^13^C enrichment were detected in metabolites of the TCA cycle (Figure 2F), suggesting that activation of NRF2 in cardiomyocytes drives changes in glycolysis and the PPP without affecting glucose contribution to the TCA cycle. To confirm these data and mitigate potential artifacts related to isolating the hearts from *in vivo* physiological conditions, we utilized an *in vivo* tail vein [U-^13^C]glucose labeling infusion strategy coupled with targeted metabolomic analysis by NMR spectroscopy as previously described^30^. In keeping with the *ex vivo* metabolomic flux results, we found a significantly increased enrichment of [2,3-^13^C] lactate, a readout for PPP flux in csKeap1KO hearts compared to WT. There were no changes between csKeap1 KO and WT hearts in forward glycolysis as measured by enrichment of [1,2,3-^13^C] lactate, pyruvate flux into the TCA cycle via oxidative decarboxylation by pyruvate dehydrogenase (PDH), or de novo glutamine synthesis from glucose^30^ (Figure 2G). Consistent with these results, we found a significant increase in gene expression of the glucose transporter *Glut1*, but not *Glut4*, and of key PPP enzymes such as glucose-6 phosphate dehydrogenase (*G6pd*), 6-Phosphogluconate dehydrogenase (*Pgd*), transaldolase (*Taldo1*), and transketolase (*Tkt*) (Figure 2D, H). Taken together, these results show that NRF2 activation in cardiomyocytes drives changes in cardiac glucose metabolism and increases glucose flux into the PPP.

**Fig. 2:**
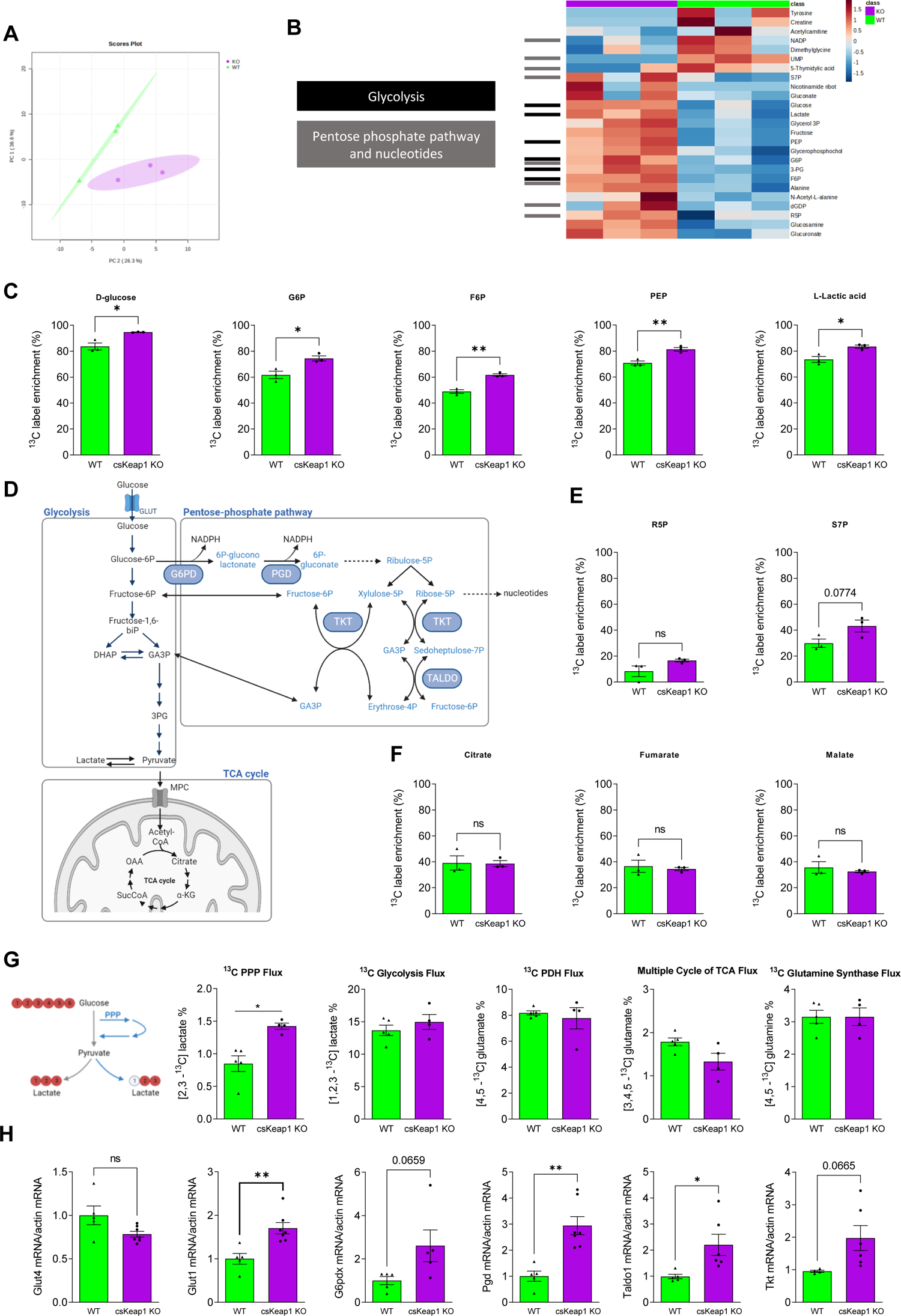
Effect of constitutive activation of NRF2 in the heart on glycolytic enzymes and ^13^C enrichment of metabolites after administration of [U-^13^C]glucose. **A)** Principal component analysis of ^13^C enrichment of metabolites following *ex vivo* Langendorff perfusion of [U-^13^C]glucose in WT and csKeap1KO mouse hearts. **B)** Heatmap showing hierarchical clustering of the top 20 ^13^C-enriched metabolites in WT vs csKeap1KO hearts. **C,E,F)** Percent ^13^C enrichment of key glycolytic pentose phosphate pathway (PPP) and tricarboxylic acid (TCA) metabolites in WT vs csKeap1KO heart following ex vivo Langendorff perfusion of [U-13C]glucose, n=3/group. **D)** Schematic representation of key metabolites of glycolysis and the PPP. **G)** Percent ^13^C enrichment of metabolites in the heart following in vivo intravenous infusion of [U-^13^C]glucose in WT and csKeap1KO mice, measured via NMR. The simplified schematic shows ^13^C labelling patterns of lactate via glycolysis or the PPP from [U-^13^C6] glucose substrate. Red fills indicate ^13^C atoms. [2,3-^13^C]lactate as a readout of PPP flux, [1,2,3-^13^C]lactate as a readout of glycolysis flux, [4,5-^13^C]glutamate as a readout of Pyruvate dehydrogenase (PDH) flux, [4,5-^13^C]glutamine as a readout of glutamine synthase flux and [3,4,5-^13^C]glutamate as a readout of multiple TCA cycle flux. n≥4/group. **H)** qPCR analysis of Glucose transporter 4 (*Glut4*), Glucose transporter 1 (*Glut1*), Glucose-6-phosphate dehydrogenase (*G6pd*), 6-Phosphogluconate dehydrogenase (*Pgd*), Transaldolase 1 (*Taldo1*) and Transketolase (*Tkt*) transcripts; n≥5/group. Data are presented as mean ± SEM. *P < 0.05, **P < 0.01, and ns, not significant by unpaired Student’s t test.

### Activation of NRF2 in cardiac-specific Keap1 knockout mice protects the heart from pressure overload-induced cardiac dysfunction

Previous studies have shown that loss of NRF2 exacerbates cardiac dysfunction and promotes heart failure in settings of stress such as pressure overload or myocardial infarction^8,10, 12^. However, whether specific activation of NRF2 in cardiac myocytes (here via Keap1KO) can protect the heart against cardiac dysfunction and failure is unclear. We therefore investigated the role of NRF2 activation under pathological conditions by subjecting csKeap1KO mice and WT littermates to suprarenal abdominal aortic constriction (AAB) or sham surgery. WT mice developed clear cardiac dysfunction compared to sham-operated mice 6 weeks after AAB surgery, as assessed by echocardiography (Figure 3A-C). Strikingly, csKeap1KO mice were protected against cardiac dysfunction as evidenced by a lack of reduction in ejection fraction and fractional shortening in comparison to sham-operated csKeap1KO mice (Figure 3A-C; Supplementary Figure 2G-H). Upon pressure overload, similar levels of cardiac hypertrophic remodeling were detected in both WT and csKeap1KO mice as evidenced by a significant increase in both heart and cardiomyocyte size, and increased mRNA expression levels of hypertrophic markers such as atrial natriuretic factor (ANF) and brain natriuretic peptide (BNP) (Figure 3D-G). However, WT hearts had higher levels of interstitial fibrosis and cardiomyocyte cell death after pressure overload-induced stress than csKeap1KO hearts, as measured by Picrosirius red and TUNEL staining, respectively (Figure 3H-J). Overall, our data showed that cardiomyocyte-specific activation of NRF2 via knock-out of Keap1 protects the heart against pressure overload-induced cardiomyocyte cell death, cardiac fibrosis, and contractile dysfunction but without significantly affecting the extent of hypertrophy.

**Fig. 3:**
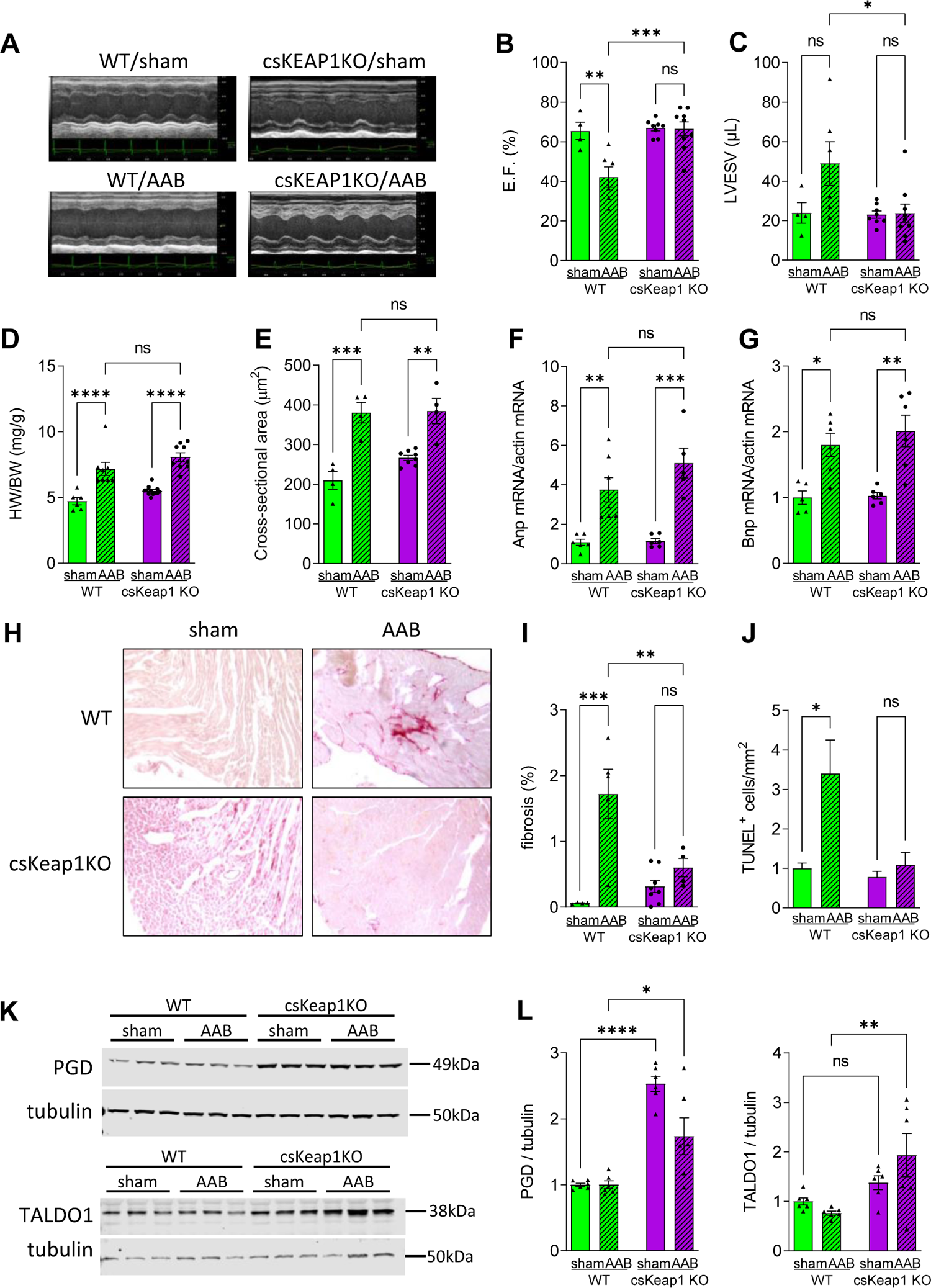
Effect of chronic pressure overload on csKeap1KO hearts. **A)** Representative M-mode echocardiography images of WT and csKeap1KO mice 6 weeks after aortic abdominal banding (AAB) or sham surgery. **B, C)** Ejection fraction (EF) and LV end-systolic volume (LVESV) in WT and csKeap1KO mice subjected to AAB or sham surgery, n≥4/group. **D)** Heart weight/body weight ratio (HW/BW), n≥6/group. **E)** Cardiomyocyte cross-sectional area in heart sections of sham or AAB-operated WT and csKeap1KO hearts, n≥4 hearts/group. **F, G)** mRNA levels of hypertrophic makers *Anp* and *Bnp*, n≥5/group. **H)** Representative images of heart sections of WT or csKeap1KO mice 6 weeks after sham or AAB surgery, stained with Picrosirius red. **I, J)** Mean data for fibrosis in heart sections from WT and csKeap1KO mice, n≥4 hearts/group. **J)** TUNEL–positive cells per square millimeter of heart tissue in sections of WT and csKeap1KO hearts; n≥3 hearts/group. Representative immunoblots **K)** and quantification data **(L)** for changes in key Pentose phosphate pathway enzymes 6-Phosphogluconate dehydrogenase (Pgd) and Transaldolase1 (Taldo1) in sham or AAB-operated WT and csKeap1KO hearts, N=6. Data are presented as means ± SEM. *P < 0.05, **P < 0.01, ***P < 0.001 and ns, not significant by two-way ANOVA, followed by Tukey’s multiple comparisons test.

We assessed how levels of PPP enzymes are changed in csKeap1KO mice in the setting of pressure overload and found upregulation at protein level of some of the PPP regulating enzymes, particularly PGD and TALDO1 in csKeap1KO hearts compared to WT littermates upon pressure overload (Figure 3K-L and Suppl. Figure 2J-K). In line with constitutive activation of NRF2, we also found that the mRNA expression levels of genes regulating antioxidant responses (*Nqo1, Gsta2, Txnrd1*), glutathione reduction (*Gsr*) and synthesis (*Gclc, Gclm*), as well as glutathione levels were increased in csKeap1KO hearts from both sham and AAB-operated mice (Suppl. Figure 2A-F, I). Overall, these data indicate that the cardioprotective effects of NRF2 activation are associated not only with an upregulation of NRF2 antioxidant targets but also NRF2-mediated glucose metabolic reprogramming and increased activation of the PPP.

### NRF2 promotes cardiomyocyte survival via activation of the PPP

To assess the contribution of NRF2-driven alterations in cardiac glucose intermediary metabolism and activation of the PPP to its protective effects in the heart, we next turned to studies in cultured cardiomyocytes. We investigated the effects of high dose isoproterenol treatment (100 µmol/L) on the viability of cultured NRVM in which the levels of Keap1 were manipulated. In line with our *in vivo* data, we found that the siRNA-mediated knockdown of Keap1 to activate NRF2 in cardiomyocytes resulted in a significant increase in expression levels of PPP enzymes including *G6pd*, *Pgd*, and *Tkt* (Figure 4A) as well as other well-established NRF2 targets (Suppl. Figure3A-B). Strikingly, we observed that activation of NRF2 by silencing of Keap1 resulted in a significant reduction in NRVM cell death under isoproterenol stimulation as compared to treatment with a scrambled siRNA control (Figure 4C). To test whether this effect was specifically mediated by activation of the PPP, NRVM were treated with 6-aminonicotinamide (6-AN, 500µmol/L), a selective inhibitor of G6PD and PGD (Figure 4B). Given that both G6PD and PGD are responsible for NADPH regeneration, we tested the efficacy of 6-AN by measuring the NADP^+^/NADPH ratio. Our results showed a clear increase in NADP^+^/NADPH levels in both control and siKeap1 transfected cells (Suppl. Figure 3C). Treatment with 6-AN did not cause a significant increase in cell death *per se* in NRVM; however, the beneficial effect of Keap1 silencing on isoproterenol-induced cell death was abolished in the presence of the PPP inhibitor 6-AN (Figure 4C). In keeping with this finding, we also found that genetic inhibition of the PPP via silencing of G6PD abolished the protective effect of NRF2 activation (Figure 4D-E). Taken together, these results indicate that NRF2-mediated activation of the PPP is necessary to prevent stress-induced cell death in cardiomyocytes.

**Fig. 4.**
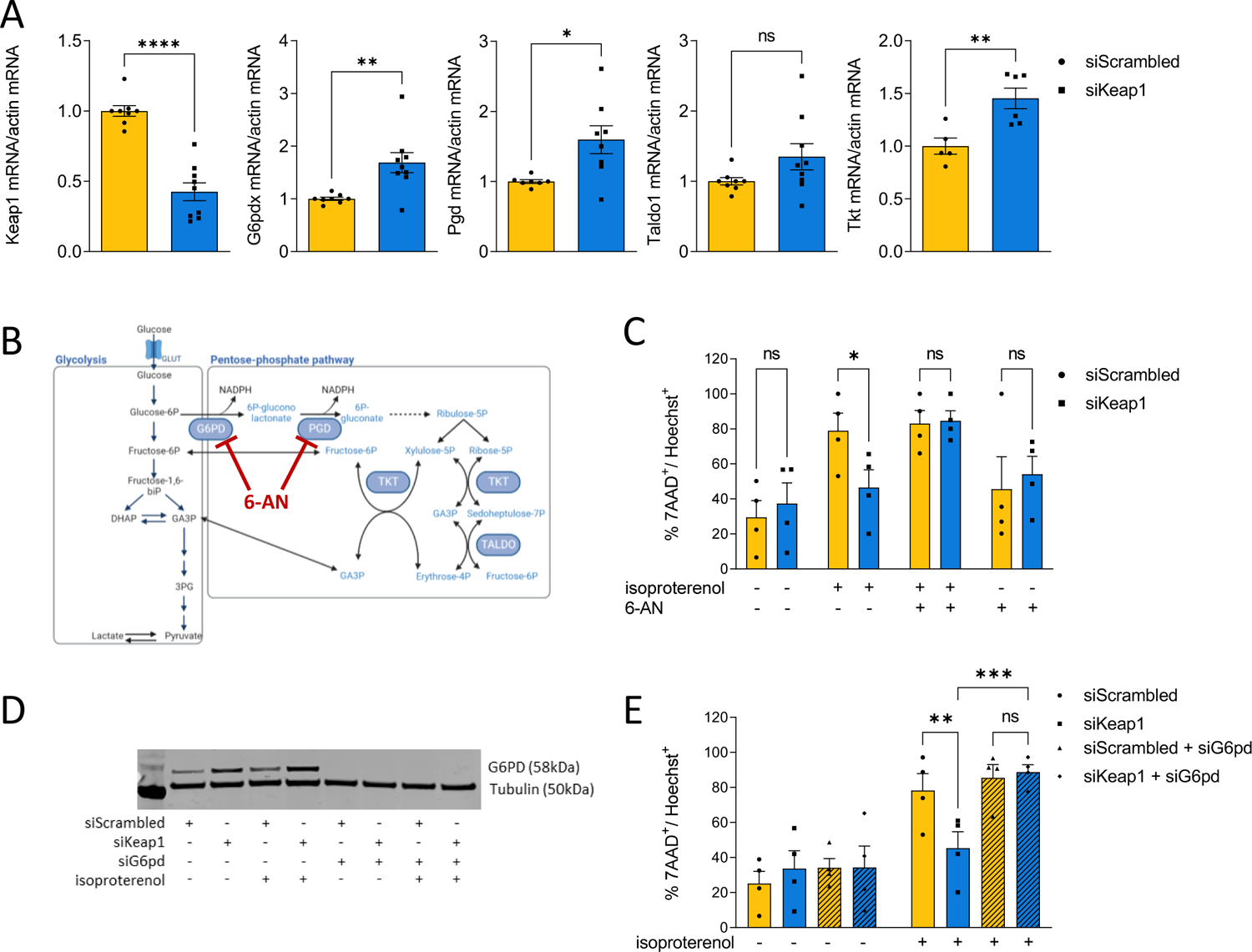
NRF2-mediated upregulation of the PPP prevents stress-induced cardiomyocyte death. **A)** mRNA levels of Keap1, Glucose-6-phosphate dehydrogenase (*G6pd*), 6-Phosphogluconate dehydrogenase (*Pgd*), Transaldolase 1 (*Taldo1*) and Transketolase (*Tkt*) measured in neonatal rat ventricular myocytes (NRVM) transfected with either control scrambled siRNA or siKeap1. n≥5/group. **B)** Schematic representation of Glycolysis and Pentose phosphate pathway (PPP) highlighting the PPP enzymes (Glucose-6-phosphate dehydrogenase, G6PD and 6-Phosphogluconate dehydrogenase, PGD) inhibited by 6-aminonicotinamide (6-AN). **C)** Cell viability measured as percentage of 7AAD-positive over Hoechst-positive cells (7AAD^+^/Hoechst^+^) in NRVM transfected with siScrambled or siKeap1, untreated or treated for 24h with different combination of isoproterenol and/or 6-AN. 4 independent experiments, n≥550 cells/group. **D)** Representative immunoblot showing efficient knockdown of Glucose-6-phosphate dehydrogenase (G6PD) in NRVM transfected with siG6pd alone or in combination with siKeap1. Tubulin was used as a loading marker. **E)** Cell viability after treatment with different combination of isoproterenol and/or 6-AN, as in panel C. 4 independent experiments, n≥600 cells/group. Data are presented as mean ± SEM. In A and C, *P < 0.05, **P < 0.01, ***P < 0.001 and ****P<0.0001 and ns, not significant by unpaired Student’s t test. In E, F and G, *P < 0.05, **P < 0.01, ***P < 0.001 and ns, not significant by two-way ANOVA, followed by Tukey’s multiple comparisons test.

### NRF2 is required to reduce stress-induced cardiomyocyte DNA damage

We next investigated the mechanism by which activation of the PPP preserves cell viability in cardiomyocytes upon neurohumoral stress. Several lines of evidence have observed DNA damage associated with stress-induced heart failure and myocardial infarction^35^. An increased level of DNA damage has been shown to exacerbate cardiomyocyte cell death and cardiac dysfunction^35, 36^, and pharmacological or genetic inhibition of DNA damage responses was found to protect the heart against pressure-overload induced cardiomyocyte hypertrophy^37^. The PPP regenerates NADPH while converting glucose-6 phosphate into ribose-5-phosphate, the sugar backbone of nucleotides. Recently, in a cell model of cancer, it was shown that the absence of the tumour suppressor p53 promotes an increase of glucose flux into the PPP which in turn stimulates more efficient DNA damage repair thus increasing cell survival upon induction of DNA damage with UV irradiation treatment^38^. In light of these findings, we assessed whether NRF2-mediated increases in PPP activity promote cardiomyocytes survival by reducing DNA damage upon stress stimuli. We first examined the changes in PPP enzymes and other NRF2 targets upon isoproterenol treatment in NRVM. Our data showed that isoproterenol treatment induced an increase in the expression level of key PPP enzymes such as *G6pd* and *Pgd*, with similar trends also observed for *Taldo1* and *Tkt* (Figure 5A). When NRF2 was silenced, these changes were abolished (Figure 5A), confirming that stress-induced gene expression of the PPP enzymes is regulated by NRF2 in cardiomyocytes. Isoproterenol treatment also increased the expression levels of other NRF2 targets such as *Nqo1* and *Gsta2*, effects which were abolished in cardiomyocytes where NRF2 was silenced (Suppl. Figure 4A-C). The treatment of cultured NRVM with isoproterenol for induced a significant decrease in cell viability as assessed by the percentage of 7-Aminoactinomycin D (7-AAD) positive NRVM. After NRF2 knockdown, isoproterenol treatment resulted in an even higher percentage of 7AAD^+^/Hoechst^+^ NRVM (Figure 5B). Of note, knockdown of NRF2 in the absence of isoproterenol also induced a decrease in cell viability *per se* in comparison to untreated control NRVM (Figure 5B). We then assessed DNA damage levels upon isoproterenol stimulation in NRVM where NRF2 was silenced. Our results showed that isoproterenol treatment induced an increase in the % of NRVM nuclei positive for the DNA-damage marker γH2AX. Importantly, we found that this effect was exacerbated in NRVM in which NRF2 was silenced (Figure 5C-D). Given that the two main products of the PPP are NADPH and the nucleotide precursor ribose-5 phosphate, we tested if exogenous supplementation of either NADPH or nucleosides was able to rescue the stress-induced DNA damage observed in NRVM lacking NRF2. Our results showed that supplementation of either NADPH or nucleosides significantly blunted the isoproterenol-induced DNA damage response in NRVM lacking NRF2 (Figure 5E-F). Overall, these data indicate that NRF2-mediated activation of the PPP is required to counteract stress-induced DNA damage responses in cardiomyocytes and that the supplementation of the PPP products, NADPH or nucleotides, reduces levels of DNA damage.

**Fig.5.**
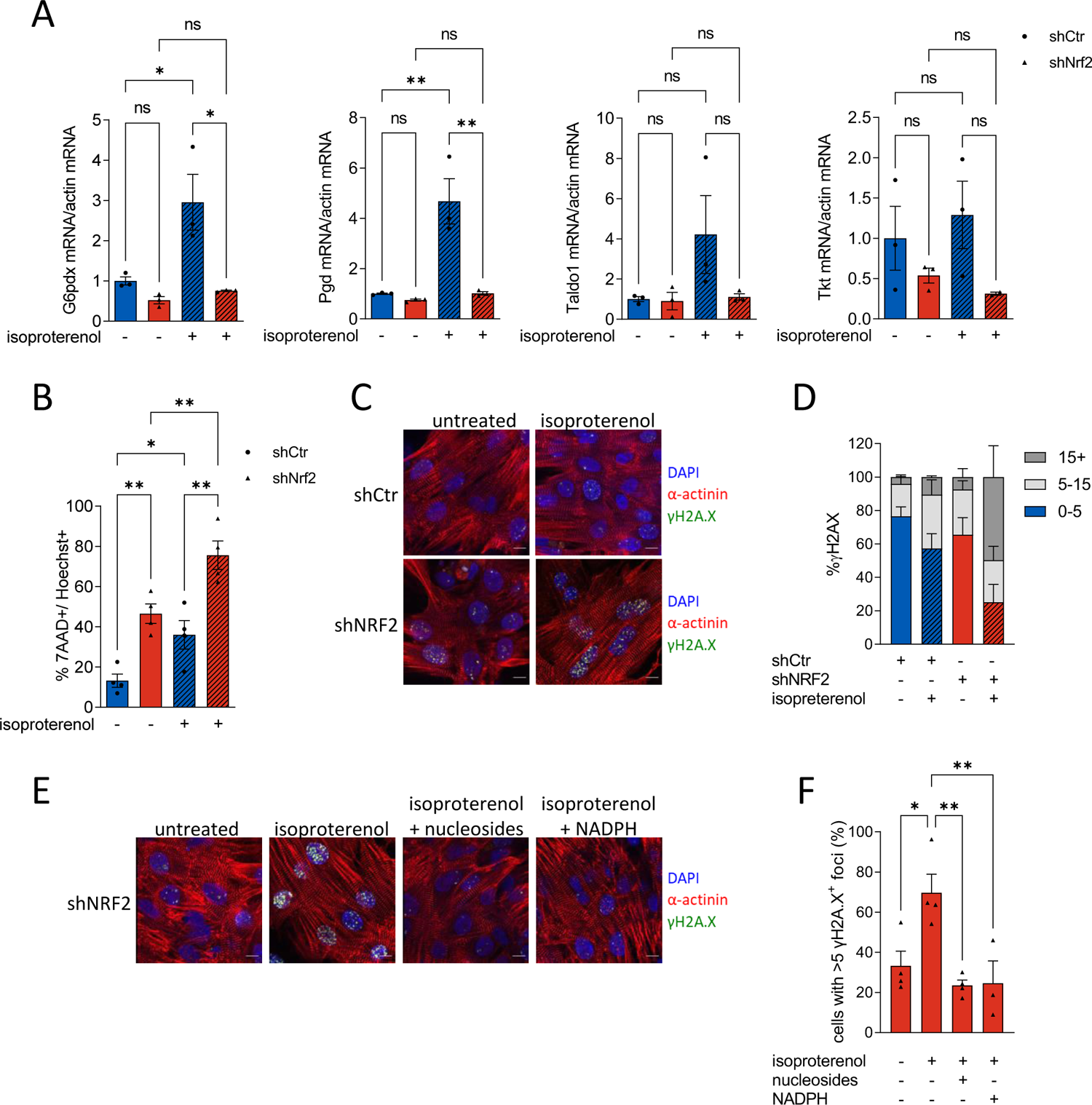
Stress-induced cardiomyocyte DNA damage is limited by NRF2 and the PPP. **A)** mRNA levels of Glucose-6-phosphate dehydrogenase (G6pd), 6-Phosphogluconate dehydrogenase (Pgd), Transaldolase 1 (Taldo1) and Transketolase (Tkt) in neonatal rat ventricular myocytes (NRVM) transduced with adenovirus expressing either control scrambled shRNA or siNrf2 and untreated or treated with isoproterenol. n=5/group. **B)** Quantification of cell viability measured as percentage of 7AAD^+^/Hoechst^+^ cells. 4 independent experiments, n≥500 cells/group. **C)** Representative immunofluorescence images of NRVM transduced with adenovirus expressing control shRNA or shNRF2 untreated or treated with 100µmol/L isoproterenol for 8 h. Cells were stained with antibodies for α-actinin, γ-H2A.X and DAPI. Scale bar: 10µm. **D)** Percentage of γ-H2A.X positive cells divided into 3 groups depending on the number of γ-H2A.X foci per nucleus: low DNA damage (0-5 foci/nucleus), moderate DNA damage (5-15 foci/ nucleus) and high DNA damage (>15 foci/nucleus). NRVM were infected with adenovirus expressing control shRNA or shNRF2, untreated or treated with isoproterenol for 8 hours. 4 independent biological replicates, n≥50 cells per condition. **E)** Representative confocal images of NRVM infected with adenovirus expressing shNRF2 and untreated or treated for 8 hours with isoproterenol alone or in combinations with nucleosides or NADPH. Cells were stained with antibodies for α-actinin, γ-H2A.X and DAPI. Scale bar: 10µm. **F)** Summary and quantification of percentage of cells with more than 5 γ-H2A.X positive foci/nucleus. NRVM were infected, treated, and stained as described in E. 3-4 biological replicates, n≥40 cells/group. Data are presented as mean ± SEM. *P < 0.05, **P < 0.01 and ns, not significant by one-way ANOVA, followed by Tukey’s multiple comparisons test.

### Activation of NRF2 protects the heart against myocardial infarction-induced DNA damage and adverse remodelling

To test whether activation of NRF2 reduces stress-induced cardiomyocyte DNA damage levels *in vivo*, we studied experimental myocardial infarction which is known to induce significant cardiomyocyte DNA damage^35^. csKeap1 KO mice and WT littermates were subjected to left anterior descending coronary artery (LAD) ligation or sham-operation. Four weeks after surgery, echocardiography data showed that csKeap1 KO mice had significantly better cardiac function than WT littermates (Figure 6A-D and Suppl. Figure 5A-C) but a similar increase in HW/BW ratio (Figure 6E). After LAD ligation, csKeap1 KO and WT littermates showed a similar infarct size and equivalent increases in cardiac fibrosis (Suppl. Figure 5D-E). When we looked at the expression levels of PPP enzymes in these experimental conditions, we confirmed an increase in mRNA and protein levels of some of the key PPP enzymes including *G6pd*, *Pdg* and *Taldo1* in sham and post-MI csKeap1 KO hearts as compared to WT littermates (Suppl. Figure 6). We performed immunohistochemistry on tissue sections from the infarcted area (border zone) or 1 mm further from the infarct area (remote zone) of csKeap1 KO hearts and WT littermates that underwent LAD permanent ligation, to assess DNA damage. The percentage of γH2AX positive cardiomyocytes was similarly high in the infarct region of csKeap1 KO and WT hearts (Figure 6F-G). However, we observed a significant reduction of MI-induced γH2AX positive cells in the remote zone of csKeap1 KO hearts compared to WT (Figure 6F-G). These data show that activation of NRF2 protects the heart against MI-induced cardiomyocyte DNA damage, adverse remodeling, and dysfunction *in vivo*.

**Fig. 6.**
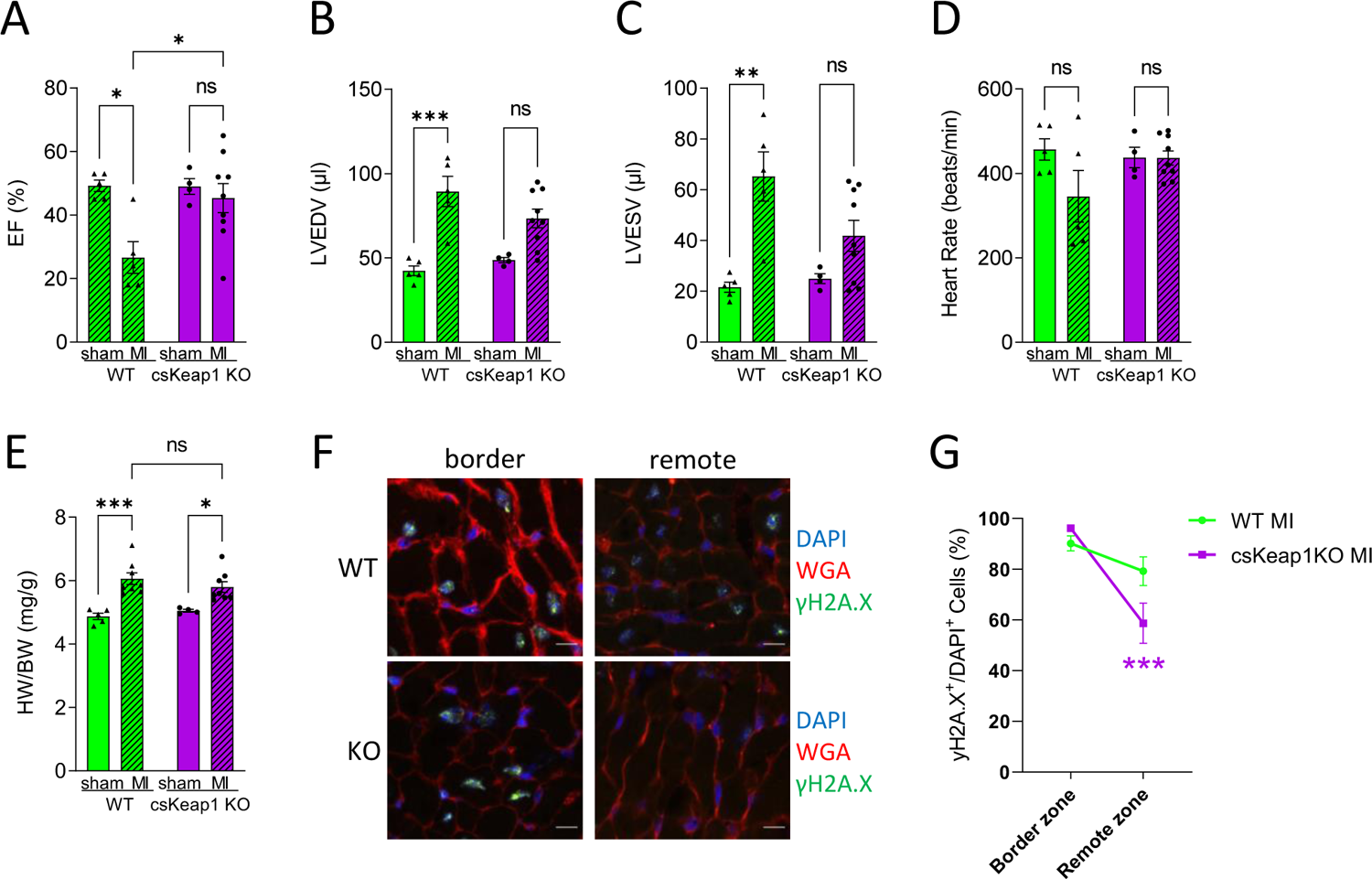
csKeap1KO hearts show better function and less DNA damage after myocardial infarction. **A-D)** Echocardiography data for ejection fraction (EF), LV end-diastolic volume (LVEDV), LV end-systolic volume (LVESV and heart rate (HR) in WT and csKeap1KO mice subjected to myocardial infarction (MI) or sham; n≥5/group. **(E)** Heart weight/body weight ratio (HW/BW) values for WT and csKeap1KO mice, 4 weeks after MI, n=8/group. **(F)** Representative confocal images of cardiac sections from remote or border zone of csKEAP1KO or WT heart after MI and stained with WGA, DAPI and for γ-H2A.X. **(G)** Percentage of γ-H2A.X positive cardiomyocytes in the border zone or remote zone; n=5 hearts/group. Data are presented as mean ± SEM. *P < 0.05, **P < 0.01, ***P < 0.001 and ns, not significant by two-way ANOVA, followed by Tukey’s multiple comparisons test.

## DISCUSSION

In this study, we identify NRF2 as a key regulator of both metabolic adaptation and antioxidant responses in the heart. Our data reveal for the first time that activation of cardiac NRF2 induces metabolic changes in the heart. We show, using complementary *in vivo* and *in vitro* approaches, that NRF2 stimulates the diversion of glucose into pathways of intermediary metabolism, in particular the PPP, which mediates cardiac protection after stress by generating both NADPH and nucleotides to prevent stress-induced DNA damage. Using state-of-the-art ^13^C_6_-glucose metabolomics stable-isotope tracing in *ex-vivo* and *in-vivo* experiments, we show that cardiac-specific activation of NRF2 leads to an increased flux of glucose carbon into the PPP but no change in glucose carbon flux into the TCA cycle. The lack of overall change in enrichment of TCA metabolites despite diversion of glucose carbons into the PPP may possibly be related to an increased expression of the glucose transporter Glut1 and increased glucose uptake, as evidenced by an increased % enrichment of ^13^C labelling of glucose in *ex-vivo* perfused csKeap1KO hearts. Thus, under these conditions, NRF2 seems to specifically regulate glucose intermediary metabolism without impacting on glucose oxidation in the mitochondria.

NRF2 is a pivotal transcription factor that regulates antioxidant responses and cell homeostasis by controlling the expression of various antioxidant, detoxification, and cytoprotective molecules. Previous studies have shown that a lack of NRF2 exacerbates pathological remodeling and cardiac dysfunction in response to a variety of disease stresses including pressure overload^8,10^, myocardial infarction^12^ and ischemia-reperfusion^39^. These cardioprotective effects of NRF2 have been attributed largely to its antioxidant and detoxification effects. Indeed, lack of NRF2 is associated with impaired antioxidant gene expression and increased oxidative stress^8,10^, a key feature of cardiac pathological remodeling and heart failure^4^. In keeping with these findings, our data also showed that cardiac-specific activation of NRF2 is associated with increased expression levels of antioxidant enzymes, both at baseline and during pressure overload, and with increased glutathione and GSH/GSSG levels. However, in other tissues, NRF2 is described to regulate not only cellular redox homeostasis but also metabolic reprogramming^14, 40–42^. For example, in lung cancer cells, NRF2 was shown to regulate the transcription of key PPP enzymes including G6PD, PGD, TALDO1 and TKT, together with other metabolic enzymes such as malic enzyme 1 (ME1) and isocitrate dehydrogenase 1 (IDH1), and genes regulating de novo nucleotide synthesis, such as phosphoribosyl pyrophosphate amidotransferase (PPAT) and methylenetetrahydrofolate dehydrogenase 2 (MTHFD2)^14^. Importantly, the NRF2 mediated expression of PPP enzymes was shown to promote diversion of glucose flux into the PPP, leading to an increase in NADPH regeneration and purine biosynthesis, and ultimately supporting tumour growth^14^. Indeed, loss-of-function mutations in the *Keap1* gene or gain-of function mutations in NRF2 gene have been associated with several types of cancer^43, 44^. Recently, loss of *Keap1* in a model of KRAS-mutant lung adenocarcinoma was shown to activate the PPP and accelerate lung tumorigenesis, while pharmacological inhibition of the PPP with 6-AN reduced tumour growth *in vivo*^40^. Overall, these recent findings describe an important new role for NRF2 as a key regulator of metabolic reprogramming and intermediary metabolism during stress^13^. Metabolic reprogramming is also a key feature of cardiac hypertrophic remodeling and heart failure. Healthy adult hearts preferentially utilize fatty acids for energy production; however, in pathological remodeling the heart decreases its capacity for fatty acid oxidation and increases glucose utilization^19–21^. Besides ATP regeneration for contraction, intermediary metabolism pathways are essential for the generation of metabolic intermediates required to enhance stress resistance (e.g., NADPH, glutathione), provide cellular building blocks (e.g., amino acids, nucleotides, lipids), and mediate cell signaling (e.g., UDP-N-Acetylglucosamine [UDP-GlcNAc], ceramides, TCA cycle intermediates). Recent evidence suggest that glucose intermediary metabolism is altered and may play a role in cardiac pathological remodeling. For example increased activity of the hexosamine biosynthesis pathway (HBP), a branch-pathway of glycolysis that generates UDP-GlcNAc which is required for the post-translational modification of proteins, has been observed in response to several cardiac stress conditions and linked to both maladaptive and adaptive outcomes^45^. Controversial results were also obtained from studies aiming at understanding the role of PPP in cardiac injury, with some evidence reporting a beneficial role of PPP activation^46, 47^ and others suggesting detrimental effects^48, 49^. It should be noted that most of these studies assessed the role of PPP only by direct modulation of the activity of glucose-6-phosphate dehydrogenase. Overall, despite an increasingly recognized role of intermediary metabolism in the heart, little knowledge is available as to how activation of cardiac glycolytic branch pathways such as the PPP influences pathological remodeling.

In this study, we unravel an important functional role of NRF2-mediated regulation of glucose intermediary metabolism via activation of the PPP. First, our *in vivo* data showed that activation of NRF2 mediates cardioprotective effects after both hemodynamic overload and myocardial infarction. In both settings, NRF2 activation protects the heart against cardiac dysfunction, fibrosis, and reduces cardiomyocyte death without influencing the extent of hypertrophy *per se*.

Second, our data reveal that NRF2 activation modulates the expression levels of PPP enzymes both *in vivo* and in primary cardiomyocytes. In csKeap1KO hearts, at baseline and upon pressure overload or myocardial infarction, we found upregulation at mRNA and protein levels of key PPP enzymes. In NRVM, we show that NRF2 is necessary to mediate isoproterenol-induced increase in expression of PPP enzymes, and that the activation of NRF2 via silencing of Keap1 silencing also induces expression of the PPP enzymes.

Third, we show that NRF2-mediated activation of the PPP is necessary to prevent stress-induced cardiomyocyte cell death and DNA damage. As such, pharmacological or genetic inhibition of the PPP abolishes the protective effects of NRF2 in NRVM. Also, activation of NRF2 and increased expression of PPP enzymes is associated with decreased level of DNA damage in infarcted heart tissues. DNA damage is associated with pathological remodeling and heart failure^35–37^, however a link between activation of the PPP and DNA damage levels has not previously been shown in cardiomyocytes or the heart. Increased PPP activity is a hallmark of many types of cancer as it plays a crucial role in providing NADPH and nucleotide precursors to sustain cancer cell survival and proliferation^50^. In a recent cancer study, p53 was shown to mediate glucose carbon diversion into the PPP, resulting in increased *de novo* nucleotide production and increased DNA damage repair and survival^38^. Similarly, in the present study we found that activation of the PPP correlates with decreased level of DNA damage upon stress stimuli in NRVM and *in vivo*. Interestingly, supplementing NRVM with either NADPH or nucleotides, the two main products of the PPP, can decrease the level of stress-induced DNA damage in cardiomyocytes where NRF2 is absent. Additional studies will be required to establish the signaling mechanisms by which supplementation with NAPDH or nucleotide reduces DNA damage in cardiac cells, and whether activation of the PPP is associated with activation of DNA damage repair pathways.

Our findings may have clinical implications as selective activation of the PPP may be considered as a novel therapeutic strategy to support adaptive effects after cardiac stress or injury. From this perspective, G6PD may represent an interesting target as recently, AG1, a small molecule that increases the activity of the wild-type, or mutant variants of G6PD have been discovered^51^. Further studies will be necessary to test the effects of this small molecule in cardiac pathological remodeling and heart failure. NRF2 is described to potentially affect several other metabolic processes, for example mitochondrial biogenesis^52, 53^, so future work is also needed to define the relative role of PPP activation versus other metabolic effects. Additionally, NRF2 may interact with transcription factors such as ATF4^54, 55^, which may itself exert metabolic effects^56, 57^. The synergistic actions of these (and other) transcription factors in mediating metabolic reprogramming in the heart therefore also merit attention.

## ACKNOWLEDGEMENTS

We thank Sharwari Verma, Shiney Reji and Mohsin Arain for the technical support, and the Wohl Cellular Imaging Centre (WCIC) at King’s College London.

---

**Supplementary Figure 1.**
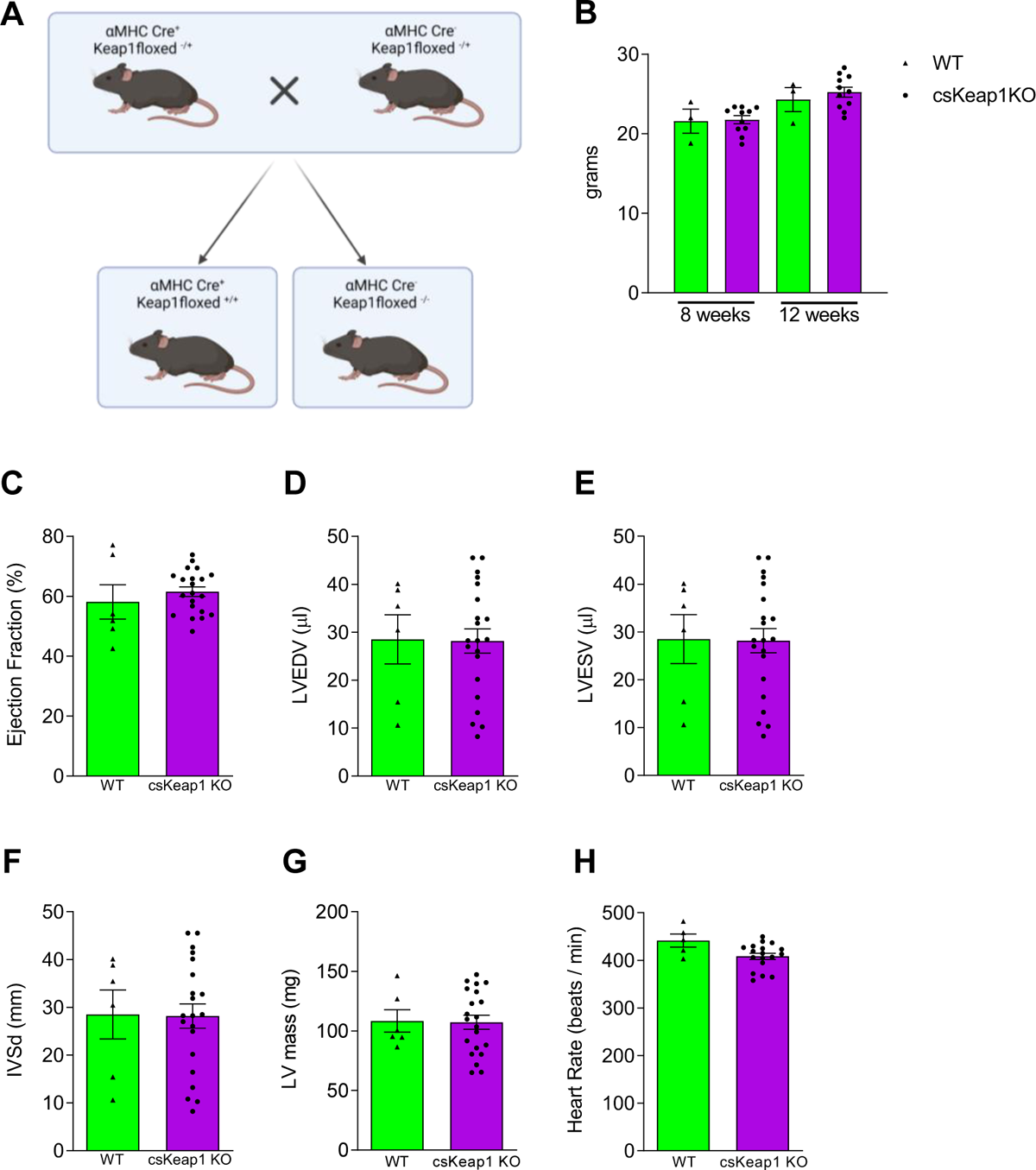
**(A)** Schematic representation of the breeding strategy. αMHC-Cre positive mice / Keap1 floxed heterozygous mice (αMHC-Cre^+^/Keap1 floxed^+/-^) were crossed with αMHC-Cre negative/Keap1 floxed heterozygous (αMHC-Cre^-^/Keap1 floxed^+/-^). αMHC-Cre^+^/Keap1 floxed^+/+^ (csKeap1KO) were compared to αMHC-Cre^-^/Keap1 floxed^-/-^ mice (WT). **(B)** Weight (g) of csKeap1KO and WT mice at 8 and 12 weeks of age, n≥3/group. **(C-H)** Echocardiography parameters and LV mass in csKeap1KO and WT littermates at 12 weeks of age. LV end-diastolic volume, LVEDV; LV end-systolic volume, LVESV; interventricular septal thickness at end diastole, IVSd. n≥6/group. Data are presented as mean ± SEM.

**Supplementary Figure 2.**
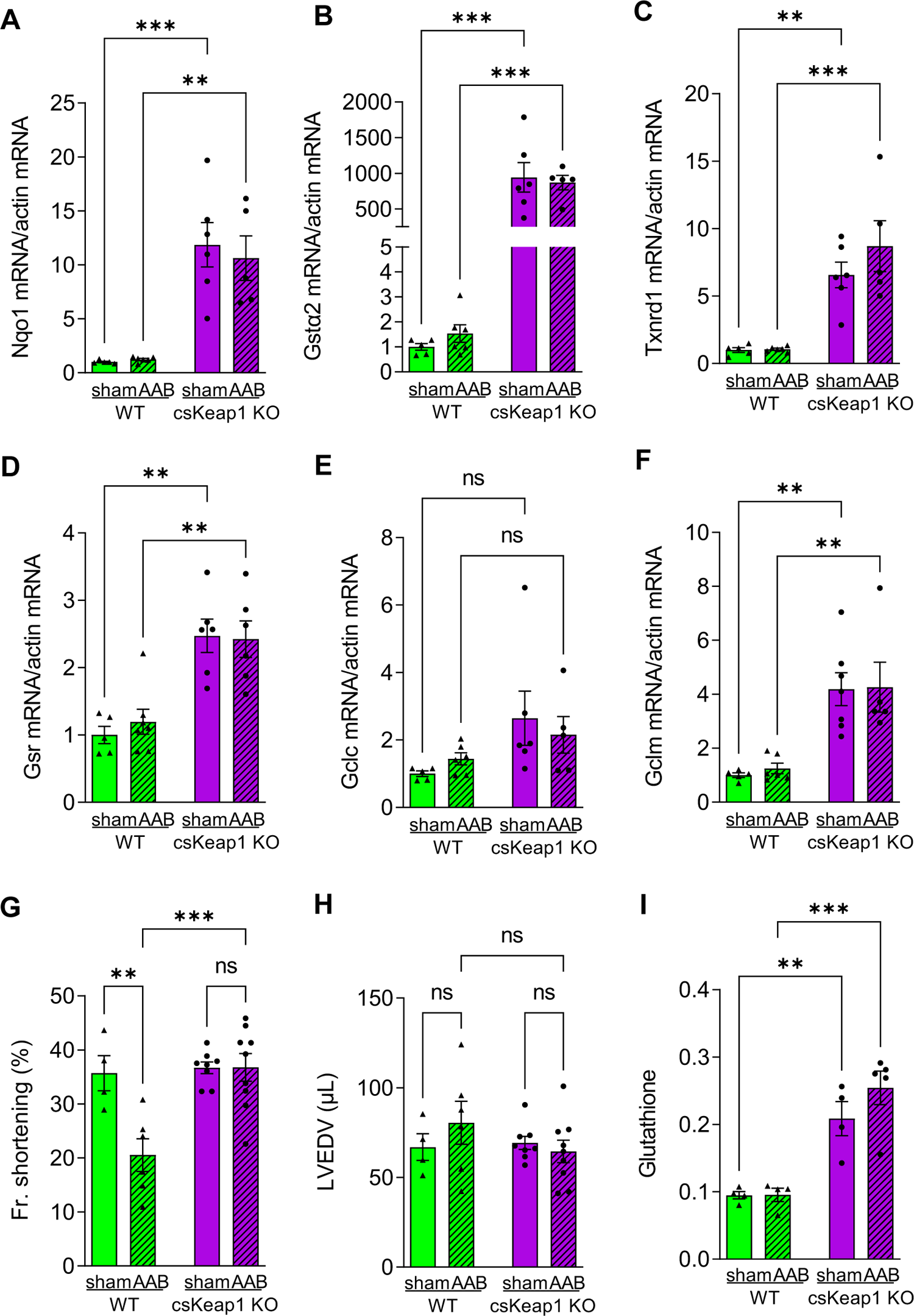
**(A-F)** mRNA levels of NRF2 targets in csKeap1KO vs WT after sham or AAB surgery; n≥4/group. *Nqo1* **(A)**, Gstα2 **(B)**, *Txnrd1* **(C)**, *Gsr* **(D)**, *Gclc* **(E)**, *Gclm* **(F)** in csKeap1KO and WT hearts upon sham or AAB surgery. **(G-H)**. Fractional (Fr.) shortening and LVEDV; n≥5/group. **(I)** Glutathione levels measured by NMR; n≥4/group. Data are presented as mean ± SEM. *P < 0.05, **P < 0.01, ***P < 0.001 and ns, not significant by two-way ANOVA, followed by Tukey’s multiple comparisons test.

**Supplementary Figure 3.**
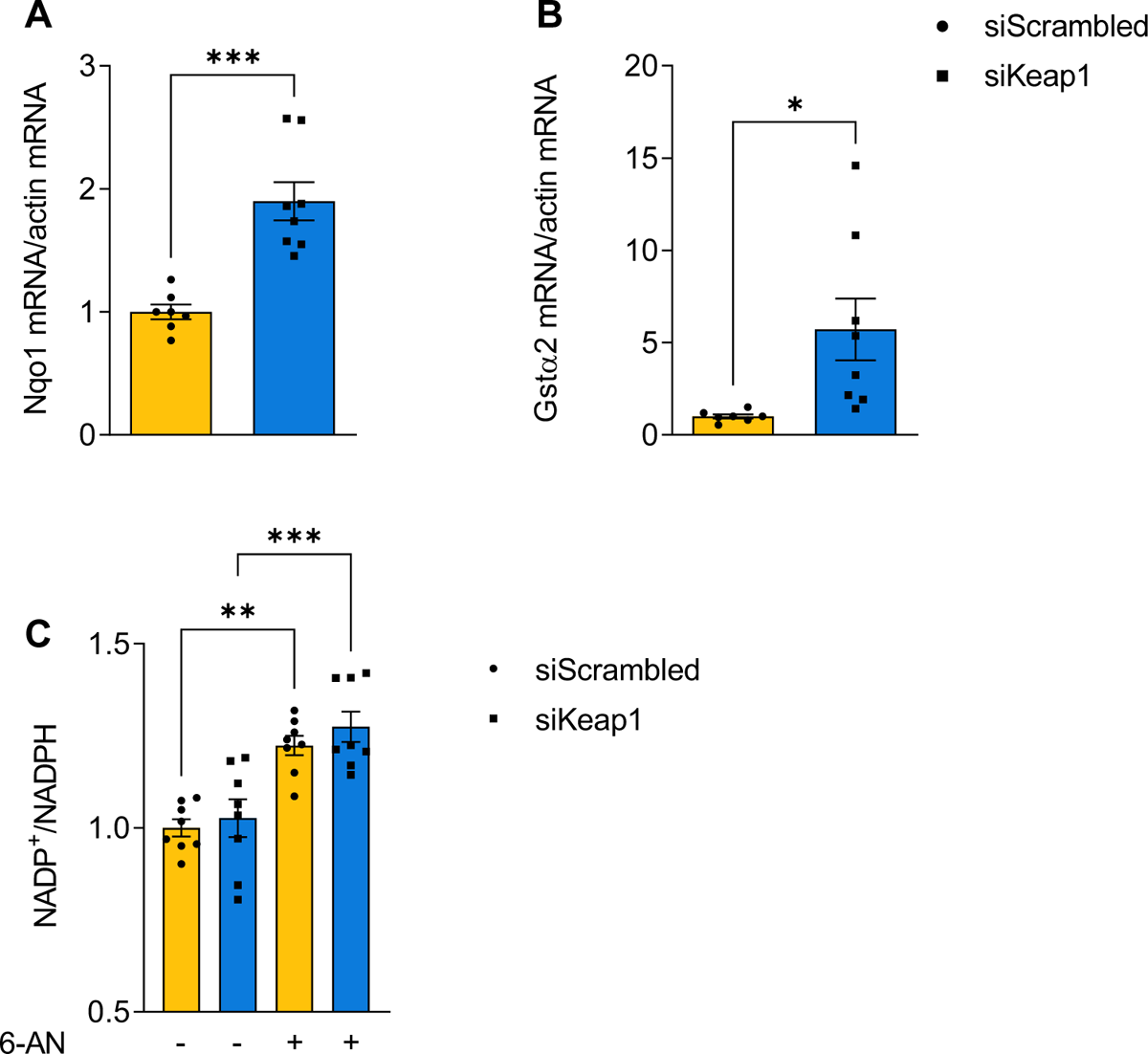
qPCR data for Nqo1. **(A)** and Gstα2 **(B)** measured in NRVM transfected with either siScrambled control or siKeap. n≥7/group. **(C)** NADP+/NADPH ratio in NRVM transfected with siScrambled or siKeap1 and treated with 2.5mmol/L 6-AN. n=8/group. Data are presented as mean ± SEM. *P < 0.05, **P < 0.01, ***P < 0.001 and ns, not significant by unpaired Student’s t test (A and B) or by two-way ANOVA, followed by Tukey’s multiple comparisons test (C).

**Supplementary Figure 4.**
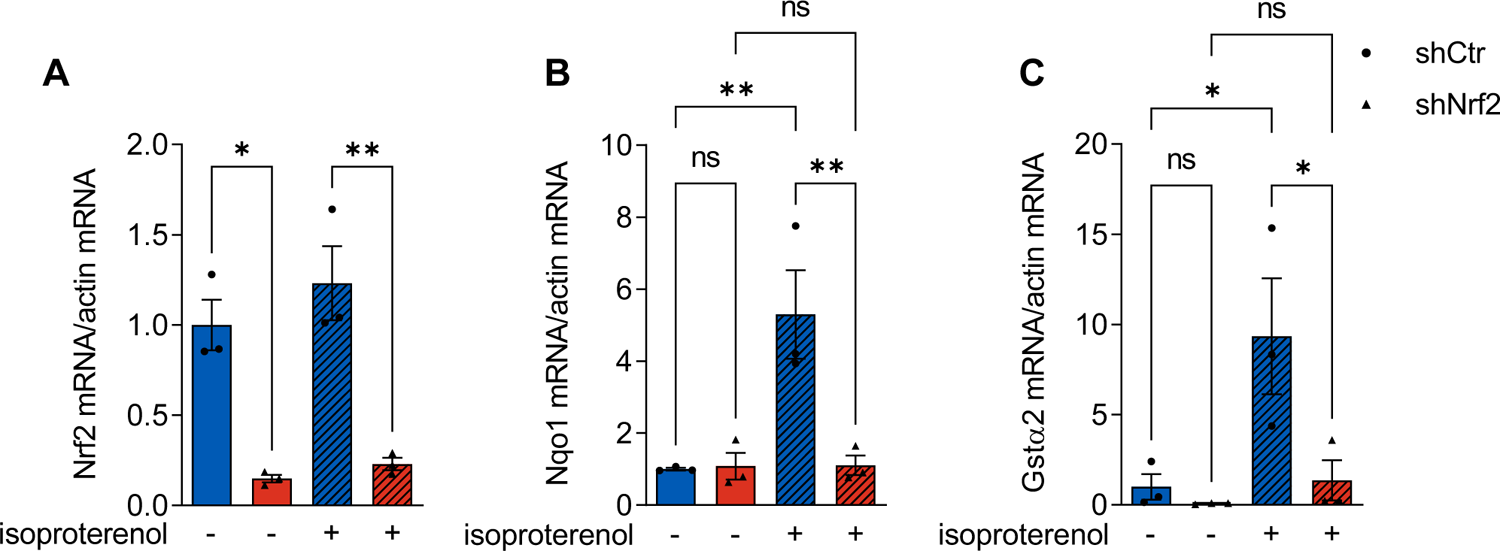
mRNA levels of *Nrf2*. **(A)**, *Nqo1* **(B)**, *Gstα2* **(C)** in NRVM infected with Ad.shCtr or Ad.shNrf2 untreated or treated with 100µmol/L isoproterenol for 24 h. N=3 biological replicates. Data are presented as mean ± SEM. *P < 0.05, **P < 0.01, ***P < 0.001 and ns, not significant by two-way ANOVA, followed by Tukey’s multiple comparisons test.

**Supplementary Figure 5.**
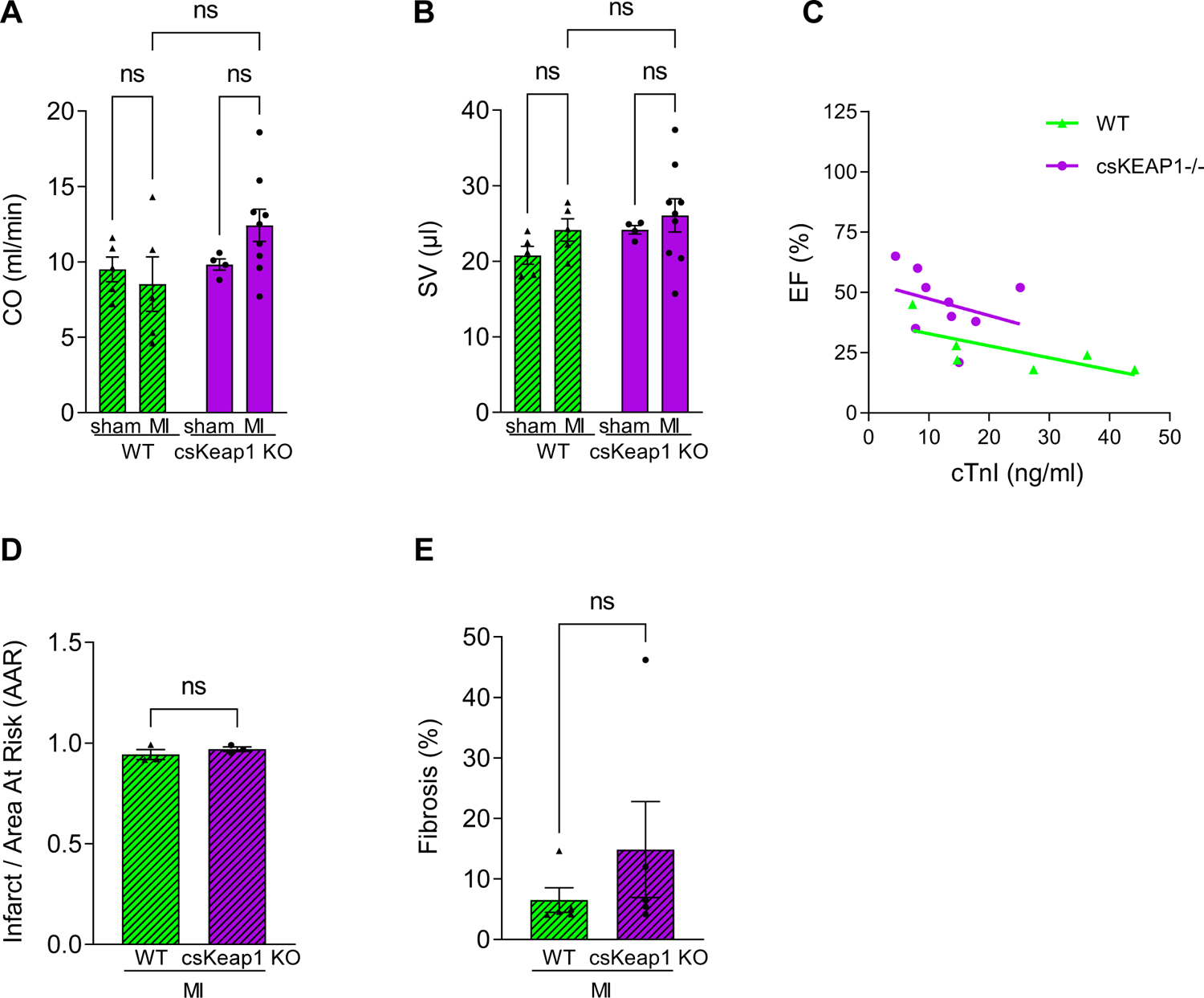
Echocardiographic parameters in csKeap1KO and WT assessed 4 weeks after sham or LAD ligation surgery; n≥7/group. Cardiac output, CO; stroke volume (SV). **(C)** EF related to infarct size as assessed by the plasma cTnI level 2 days after MI. n=6-8/group. For any given infarct size, EF is higher in csKeap1KO mice. **(D)** Infarct size over area at risk assessed by Evans blue staining in WT and csKeap1KO hearts. n=3/group. **(E)** % fibrosis assessed by Picrosirius red staining of heart sections 4 weeks after sham or LAD ligation surgery. n=5/group. Data are presented as mean ± SEM. *P < 0.05, **P < 0.01, ***P < 0.001 and ns, not significant by two-way ANOVA, followed by Tukey’s multiple comparisons test (A-D) or by unpaired Student’s t test (E-F).

**Supplementary Figure 6.**
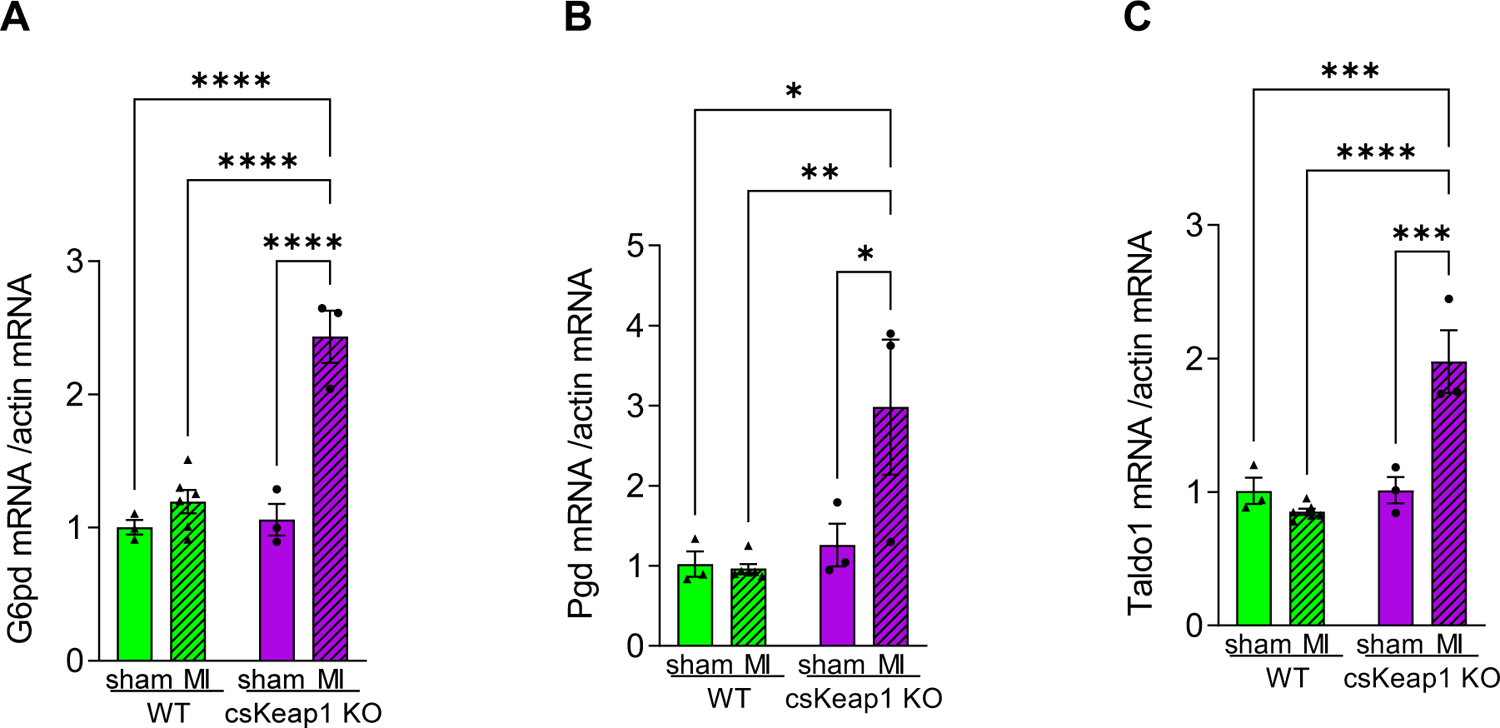
mRNA levels of G6pd. **(A)**, Pgd **(B)** and Taldo1 **(C)** in WT or csKeap1KO hearts 4 weeks after sham or LAD ligation surgery. n≥3/group. *P < 0.05, **P < 0.01, ***P < 0.001, ****P<0.0001 and ns, not significant by two-way ANOVA, followed by Tukey’s multiple comparisons test.

